# In vitro evaluation of factor IX as novel treatment for factor XI deficiency

**DOI:** 10.1101/614180

**Authors:** Kamran Bakhtiari, Joost C.M. Meijers

## Abstract

Factor XI deficiency is associated with mild to moderate bleeding upon injury. Treatment of bleeding in patients can be a challenge due to the limited availability of factor XI concentrates that may also have thrombotic side effects, and the volume overload as a result of plasma transfusion. In our in vitro study, we established that recombinant factor IX concentrate Benefix® (Pfizer) was able to potently enhance thrombin generation in factor XI depleted plasma when coagulation was initiated via the extrinsic pathway. This was due to the contamination of Benefix® with very low amounts of factor IXa that compensated for the lack of factor XI in plasma. Our data suggest that bleeding due to factor XI deficiency or antithrombotic therapy targeting factor XI may be treated with certain factor IX concentrates, which should be investigated in future clinical studies.

## INTRODUCTION

Factor XI is a zymogen of a serine protease that participates in the intrinsic pathway of coagulation. Deficiency of factor XI, first described in 1953,^1^ is characterized by mild to moderate bleeding after injury in tissues with high fibrinolytic activity such as the oral and nasal cavities and the urinary tract.^2^ Spontaneous bleeding is rare in factor XI deficient patients. Treatment of factor XI deficiency is a challenge because administration of fresh frozen plasma may lead to volume overload, and factor XI concentrates are not widely available. There are two plasma-derived concentrates, Hemoleven (a high purity product from LFB Biomedicaments, France) and FXI Concentrate (a partially purified product from Bioproducts Laboratory, UK), that have shown efficacy, but also demonstrated safety concerns.^3,4^ Traces of factor XIa in the concentrates were thought to lead to thrombosis and disseminated intravascular coagulation, and current products contain heparin, antithrombin and C1 inhibitor to eliminate factor XIa enzymatic activity. Also, other products have been used, such as recombinant factor VIIa with or without antifibrinolytic agents, but it is not clear which therapy is currently the best option.^2^ We investigated the effect of coagulation protein supplementation on thrombin generation in factor XI depleted plasma and observed a procoagulant effect of factor IX.

## METHODS

Thrombin generation in platelet-poor plasma was measured with the Calibrated Automated Thrombogram (Thrombinoscope BV, Maastricht, the Netherlands) as previously described.^5^ Briefly, 80 µl plasma was added to 20 µl of trigger reagent and incubated for 10 min at 37°C. Thrombin generation was initiated by adding 20 µl Hepes/BSA buffer containing 100 mM CaCl_2_ and 2.5 mM fluorogenic substrate Z-Gly-Gly-Arg-AMC (Bachem, Bubendorf, Switzerland). Tissue factor-dependent thrombin generation was assessed with PPP Reagent low (1 pM tissue factor in the presence of 4 µM phospholipids. Fluorescence was monitored using the Fluoroskan Ascent fluorometer (ThermoLabsystems, Helsinki, Finland), and the lag time, peak thrombin, velocity index and area under the curve (ETP) were calculated using the Thrombinoscope® software (Thrombinoscope BV). Normal pooled plasma (obtained from more than 200 healthy individuals) or factor XI depleted plasma (Siemens Healthcare Diagnostics, Marburg, Germany) were supplemented with prothrombin, factor X (both generously provided by the late dr. W. Kisiel, University of Albuquerque, NM), factor VIII (Refacto®, Pfizer, New York, NY), factor V (Haematologic Technologies Inc., Essex Junction, VT), recombinant factor IX (Benefix®, Pfizer), plasma-derived factor IX (Nonafact®, Sanquin, Amsterdam, the Netherlands), activated factor IX^6^, inhibitory antibodies against factor IX (IEP-5F5, prepared in our laboratory) and/or inhibitory antibodies against factor XI^7^ (mix of αFXI-175 and αFXI-203).

Factor IXa in factor IX preparations was determined with the Rossix Factor IXa kit (Mölndal, Sweden).

## RESULTS AND DISCUSSION

Factor XI contributes to thrombin generation when coagulation is initiated with low amounts of tissue factor.^8^ Therefore, thrombin generation initiated with 1 pM tissue factor was determined in factor XI depleted plasma supplemented with downstream coagulation proteins at their respective plasma concentrations (Figure 1A). Minor procoagulant effects on thrombin generation were observed for prothrombin (increased endogenous thrombin potential) and factor X (reduced lag time). Addition of factor VIII had no effect, while factor V reduced thrombin generation. Strikingly, a strong procoagulant effect was observed with addition of recombinant factor IX (Benefix®, Pfizer). This enhancing factor IX effect was dependent on factor IX itself, as inhibitory anti-factor IX antibodies completely abolished the procoagulant effects of recombinant factor IX (Figure 1B). The procoagulant effect was dose-dependent as increasing concentrations recombinant factor IX considerably enhanced thrombin generation in factor XI depleted plasma (Figure 1C). We evaluated the effects of a commercial plasma-derived factor IX (Nonafact®, Sanquin), but that only had a minor effect on thrombin generation in factor XI depleted plasma (Figure 1D). A major difference between the two preparations is the concentration of contaminating factor IXa. Nonafact® does not contain detectable amounts of factor IXa (<0.03% factor IXa), while Benefix® contains 0.06-0.08% factor IXa. Although these factor IXa concentrations in Benefix® are very low, they are sufficient to enhance thrombin generation and compensate for the absence of factor XI upon activation of the extrinsic pathway (Figure 2A), Finally, we tested if recombinant factor IX could also improve thrombin generation in the presence of inhibiting factor XI antibodies to mimic the situation in a factor XI deficient patient with inhibitors. Addition of anti-factor XI antibodies reduced thrombin generation in normal plasma (Figure 2B). As expected, the procoagulants effect of Benefix® were not affected by anti-factor XI antibodies, both in plasma containing factor XI (Figure 2B) and factor XI depleted plasma (Figure 2C).

**Figure 1:**
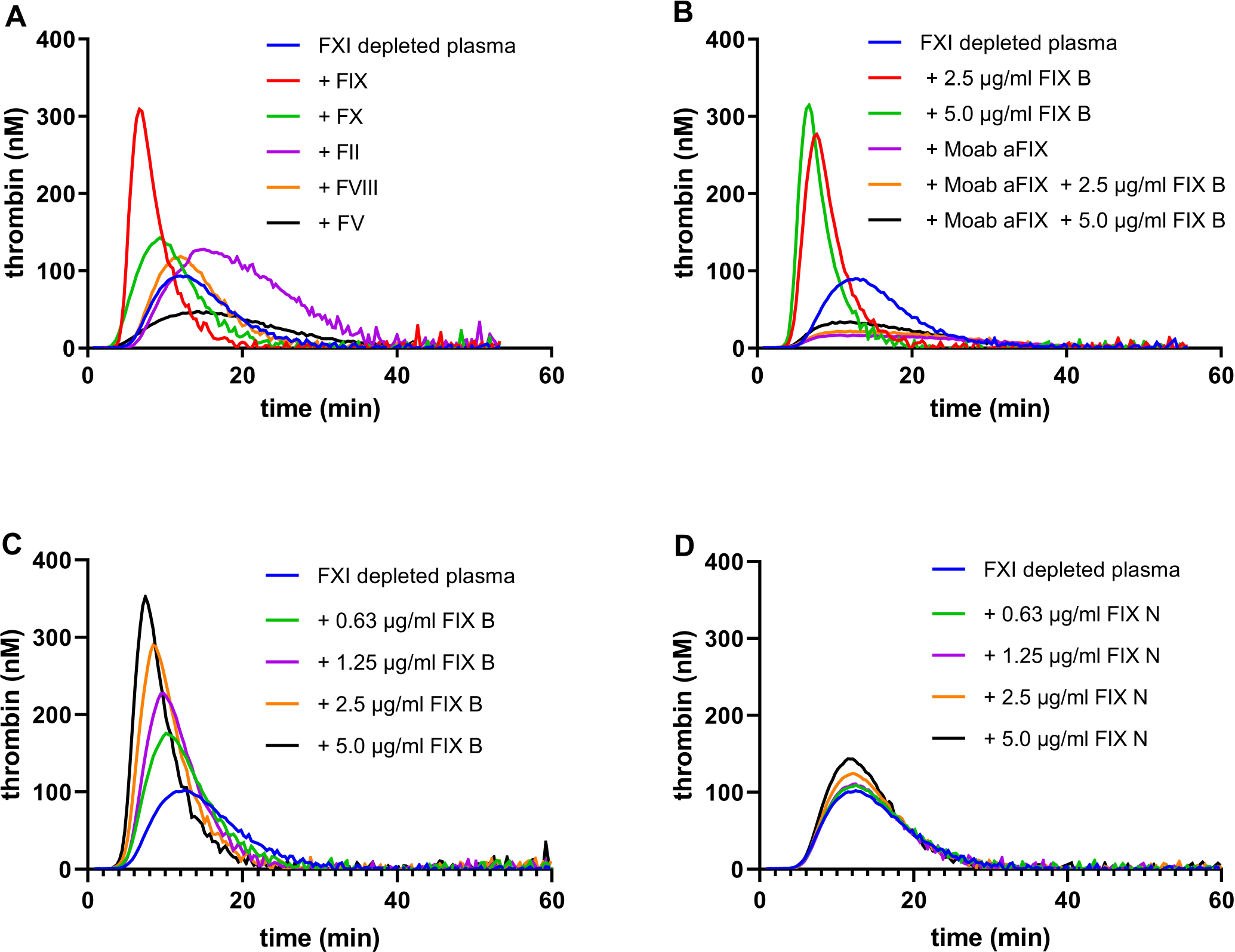
Factor IX enhances thrombin generation in factor XI depleted plasma. **A.** Factor XI depleted plasma was supplemented with 1 U/ml prothrombin (90 µg/ml), factor X (8 µg/ml), factor V (10 µg/ml), factor VIII, or factor IX (Benefix®, 5 µg/ml). Thrombin generation was initiated with 1 pM tissue factor. **B.** Factor XI depleted plasma was supplemented with 2.5 or 5 µg/ml factor IX (FIX B, Benefix®) in the presence or absence of a blocking anti-factor IX antibody (Moab aFIX). Thrombin generation was initiated with 1 pM tissue factor. **C.** Factor XI depleted plasma was supplemented with factor IX (FIX B, Benefix®) at the indicated concentrations. Thrombin generation was initiated with 1 pM tissue factor. **D.** Factor XI depleted plasma was supplemented with factor IX (FIX N, Nonafact®) at the indicated concentrations. Thrombin generation was initiated with 1 pM tissue factor.

**Figure 2:**
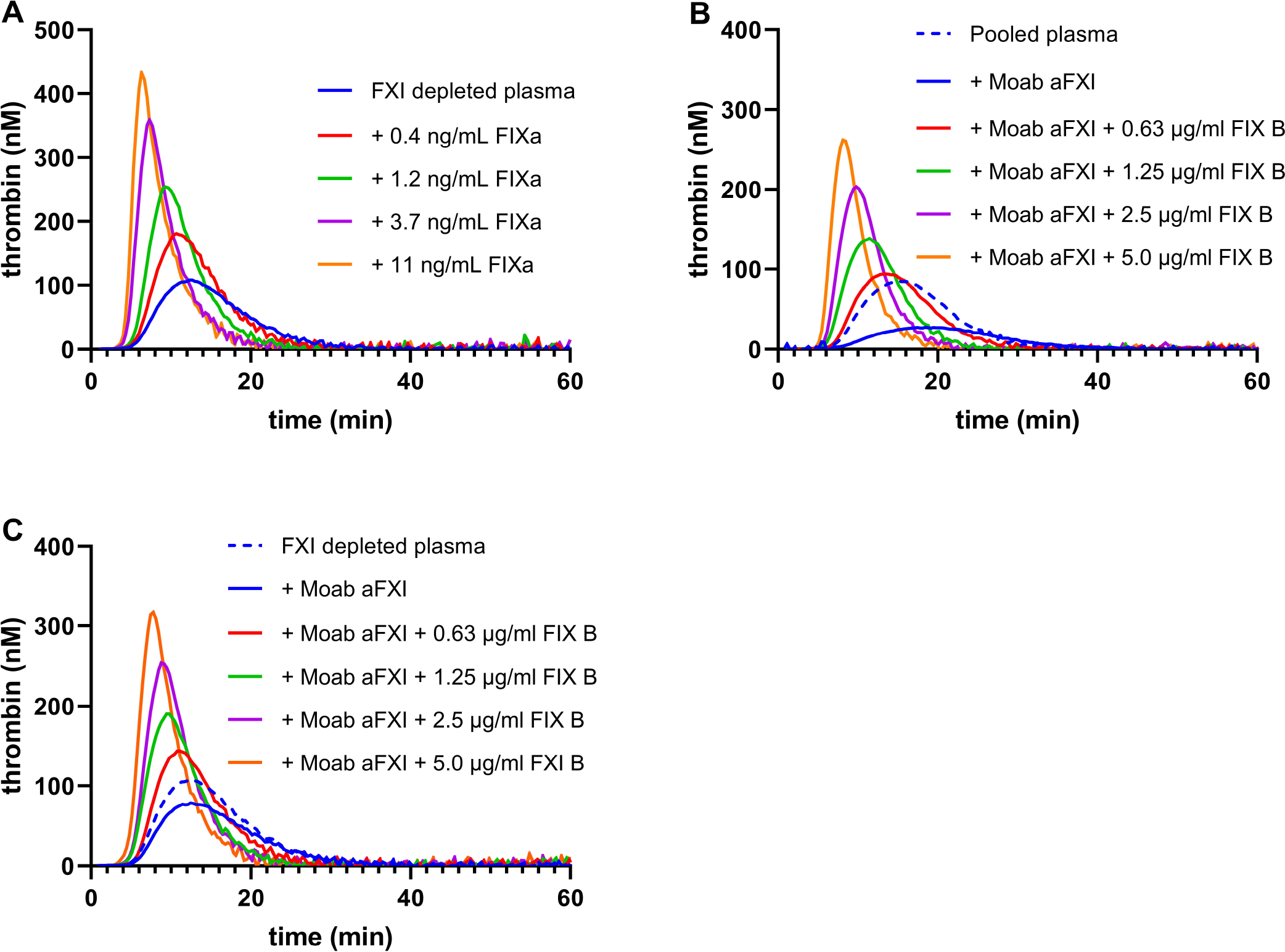
Procoagulant effect of factor IX concentrate is caused by contamination with factor IXa. **A.** Factor XI depleted plasma was supplemented with factor IXa at the indicated concentrations. Thrombin generation was initiated with 1 pM tissue factor. **B.** Normal pooled plasma was supplemented with recombinant factor IX (FIX B, Benefix®) at the indicated concentrations in the presence of inhibitory anti-factor XI antibodies (Moab aFXI). Thrombin generation was initiated with 1 pM tissue factor. **C.** Factor XI depleted plasma was supplemented with recombinant factor IX (FIX B, Benefix®) at the indicated concentrations in the presence of inhibitory anti-factor XI antibodies (Moab aFXI). Thrombin generation was initiated with 1 pM tissue factor.

Our data may have important implications for the treatment of factor XI deficient patients. Since Benefix® is a registered drug and is used in many hemophilia B patients, there is considerable evidence for its safety. Most likely, other factor IX products will also be able to improve coagulation in factor XI deficiency, but that will be dependent on their contamination with factor IXa. This can be easily determined using the methodology of our study. Furthermore, it appears that the dose of factor IX for factor XI deficiency can be lower than for factor IX deficiency, since supplementation with 0.63 µg/ml (~0.13 U/ml) Benefix® already provided considerable additional thrombin generation in factor XI depleted plasma (Figure 1C). Clinical studies are urgently needed to establish if factor IX products can be used for treatment of bleeding in factor XI deficient patients. In addition, since factor XI is an interesting novel target for antithrombotic therapy with low bleeding risk,^9,10^ the use of factor IX concentrates may be considered in those patients treated with anti-factor XI therapy that still develop bleedings.

## Acknowledgments

### Authorship Contributions

KB: designed and performed research, analyzed data, approved manuscript JCMM: concept, design and supervision of research, analyzed data, wrote the manuscript

### Disclosure of Conflict of Interest

The authors are employed by Sanquin, the manufacturer of Nonafact®. The employer of JCMM (Sanquin) received honoraria for participation in scientific advisory board panels and consulting for Bayer and Daiichi Sankyo.

